# Stimulus-specific adaptation to behaviorally-relevant sounds in awake rats

**DOI:** 10.1101/732826

**Authors:** A. Yaron, M. M. Jankowski, R. Badrieh, I. Nelken

**Author notes:** Corresponding author: Israel Nelken, The Edmond and Lily Safra Center for Brain Sciences, Edmond J. Safra Campus, Jerusalem 91904, Israel.

## Abstract

Stimulus-specific adaptation (SSA) is the reduction in responses to a common stimulus that does not generalize, or only partially generalizes, to other stimuli. SSA has been studied mainly with sounds that bear no behavioral meaning. We hypothesized that the acquisition of behavioral meaning by a sound should modify the amount of SSA evoked by that sound. To test this hypothesis, we used fear conditioning in rats, using two word-like stimuli, derived from the English words “danger” and “safety”, as well as pure tones. One stimulus (CS+) was associated with a foot shock whereas the other stimulus (CS-) was presented without a concomitant foot shock. We recorded neural responses to the auditory stimuli using chronically implanted multi-electrode arrays, recording responses telemetrically in freely moving animals before and after conditioning. Consistent with our hypothesis, SSA changed in a way that depended on the behavioral role of the sound: the contrast between standard and deviant responses remained the same or decreased for CS+ stimuli but increased for CS- stimuli, showing that SSA is shaped by experience. In most cases the sensory responses underlying these changes in SSA increased following conditioning. Unexpectedly, the responses to CS+ word-like stimuli showed a specific, substantial decrease, which we interpret as evidence for substantial inhibitory plasticity.

## Introduction

Neural responses throughout the auditory system show sensitivity to stimulus probability. Such sensitivity is often probed using oddball sequences (1). In an oddball sequence, a common (standard) sound and a rare (deviant) sound are randomly intermixed. The concomitant reduction in the response to the common stimulus that does not generalize, or only partially generalizes, to other, rare stimuli, was named stimulus specific adaptation, SSA (2). SSA has been demonstrated in the auditory system of many mammalian species, including cats, rats, mice, gerbils, macaques, and bats (1,3–10) as well as in birds (11–14). In addition to auditory cortex, SSA (at least for pure tones) has been found in rat inferior colliculus (8,15,16), rat thalamic reticular nucleus (17), and the medial geniculate body (MGB) of rats (5) and mice (3), but not in the in the rat cochlear nucleus (18). Most studies of SSA used pure tones of different frequencies as standards and deviants. More recently, we demonstrated SSA for complex sounds (19). In particular, we demonstrated SSA for word-like stimuli that have been acoustically adapted to the rat auditory system.

Previous studies of SSA have used sounds that did not carry a behavioral meaning. The current study was designed to explore how the behavioral relevance of sounds affects the probability dependence of the responses they evoke. Functionally, it may be advantageous to reduce the adaptation of the responses evoked by a sound which predicts a negative consequence (e.g. CS+ sounds in discriminative fear conditioning paradigms), in order to ensure a robust neuronal representation of such sounds. Such a change would make the responses to standards and deviants more similar to each other, and SSA for such a sound would become smaller. The reverse may be advantageous for sounds that are associated with a neutral consequence (e.g. CS- sounds in discriminative fear conditioning paradigms): by the same stronger adaptation and therefore larger SSA following conditioning.

It is now well established that learning modifies systematically the representation of acoustic information in A1. Shifts of frequency tuning that favor behaviorally important frequencies are a consistent finding across many types of training, reinforcement motivation, and laboratories. Plasticity in A1 underlies at least some features of auditory memory (20,21). Fear conditioning is an easy and robust way of modifying animal behavior (22). When used with pure tones, the plastic changes that fear conditioning induces in the auditory system are reasonably well-understood (23–25). We therefore used fear conditioning to explore the interaction of learning with SSA.

We used both pure tones and the word-like stimuli developed in Nelken et al. (19) for discriminative fear conditioning, and measured the SSA evoked by these sounds before and after conditioning. SSA indeed tended to decrease for the CS+ and increase for the CS- sounds following conditioning. Unexpectedly, the patterns of changes in the neural responses that led to these consequences was dependent on the acoustic structure of the stimuli used during conditioning. Consistent with previous findings, conditioning with pure tones increased neural responses to all stimuli. In contrast, conditioning with word-like stimuli led to a specific and surprisingly large decrease in the responses to the CS+ stimulus.

## Materials and Methods

### Animals

The joint ethics committee (IACUC) of the Hebrew University and Hadassah Medical Center approved the study protocol for animal welfare. The Hebrew University is an AAALAC International accredited institute. We used 21 adult female Sabra rats for this study (Harlan Laboratories Ltd., Jerusalem, Israel). The rats were kept in a temperature and humidity-controlled room, maintained on a 12-h light/dark cycle (lights on from 07:00 to 19:00), and had free access to water and standard rodent food pellets (Harlan Laboratories) except during the recording sessions.

### Experimental Design

The timeline of the experiment is described in Fig 1. On week 1, rats were habituated to handling for 5 days, 20 min each day. On week 2 rats were habituated to the experimental cage (a 53×35 cm box with a grid floor, Med Associates, Inc.; context A), 20 min each day. On day 15, the rats went through electrode implantation surgery and left to recover for 3 days. Responses to auditory stimuli (see below for details) were collected for 2 days and then, to confirm stability of the recordings, for another 2 days a week later. Five days after the conclusion of the recording sessions, the rats underwent conditioning. One and two days following conditioning, the rats were tested for freezing in a different context (context B) and auditory responses were collected again. Context B had a black plastic floor placed over the metal grid floor, and a blue plastic sheet was placed around the walls, modifying the shape of the box. The conditioning and test boxes and the grid floor were cleaned before and after each session with 70% ethanol.

**Fig 1.**
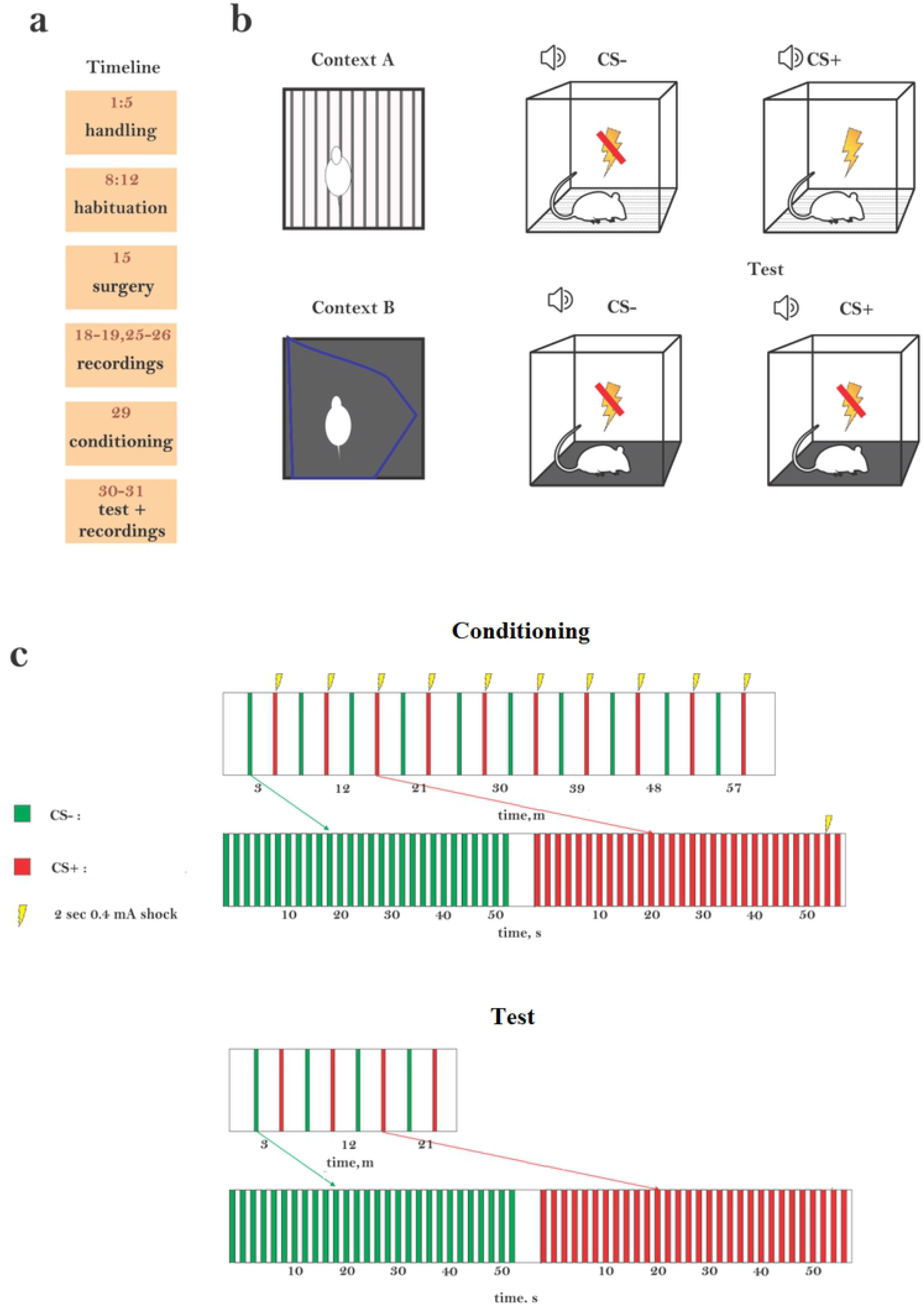
Experimental design. (a) The timeline of the experiment (in days). (b) Conditioning was performed in context A (CS+ coupled with foot shocks), testing was performed in context B (CS+ without foot shocks). (c) To induce conditioning, animals were exposed to 20 blocks of sounds, alternating between CS- (green) and CS+ (red). A block consisted of a 30s train of one of the stimuli delivered at 0.5 Hz. CS+ was paired with a foot shock (2 s, 0.4 mA). The onset of the foot shock was 2 seconds before the end of the sequence. In a fear retrieval test, rats received alternately 4 presentations of CS– and 4 presentations of CS+ stimuli with no shock associated with either.

### Surgical procedure

Rats were anesthetized initially in an induction chamber with sevoflurane (8% in oxygen, Piramal Critical Care Inc., Bethlehem, PA, USA). Their heads were shaved and they were placed in a stereotaxic instrument with a mask for gas anesthesia (David Kopf Instruments, CA, USA). Sevoflurane concentration was slowly adjusted to the level of 2-2.5% and maintained at this level throughout the surgery. Surgical level of anesthesia was verified by lack of pedal-withdrawal reflex. The eyes were protected with a thick layer of vaseline and the skin on the head was disinfected with povidone-iodine solution (10%, equivalent of 1% iodine, Rekah Pharm. Ind. Ltd., Holon, Israel).

A 1.5-2 cm longitudinal cut of the skin on the head was made and the bones of the skull were exposed. The connective tissue was mechanically removed from the skull and bones were treated with a 15% hydrogen peroxide solution (Sigma Aldrich Inc., St. Louis, MO, USA) which was immediately flushed with sterile saline. When the surface of the skull was clean and dry, a reference point for the implantation of recording electrodes was marked. Subsequently, 7-8 holes for supporting screws were drilled and screws were mounted in the skull. The screws were fixed together and to the bone with dental cement (Coral-fix, Tel Aviv, Israel) forming a base for the implant. The electrode implantation site was kept free of dental cement.

A small opening was drilled in the skull above auditory cortex and the dura was removed. Rats were implanted with custom designed 16 electrode arrays (MEA, Microprobes for Life Sciences, Gaithersburg, MD). The electrodes were 75-micron diameter Parylene C coated tungsten wires with a nominal impedance of 1MΩ. Beyond the epoxy, their length was 4 mm. They were organized in a 4X4 square with 0.3 mm spacing.

The electrodes were implanted using a stereotaxic Instrument (David Kopf Instruments, Tujunga, California), vertically, just medial to the lateral ridge, at coordinates targeted to the left primary auditory cortex (5 mm posterior to bregma, 2.3-2.4 mm below brain surface). While lowering of the electrodes inside the brain, responses to auditory stimuli were recorded and the final depth of the electrodes was set accordingly. The array was fixed to the base of dental cement previously prepared on the skull. The ground wire was soldered to one of the screws and insulated.

The wounds were cleaned and treated in situ with antibiotic ointment (synthomycine, chloramfenicol 5%, Rekah Pharm. Ind. Ltd., Holon, Israel) and dermatol (bismuthisubgallate, Floris, Kiryat Bialik, Israel). To prevent postoperative pain, rats received subcutaneous injection of Carprofen 50 mg/ml (5% W/V) in a dose of about 13 mg/kg (Norocarp, Norbrook Laboratories Limited, Newry, Co. Down, Northern Ireland) immediately following the surgery. Injections of Carprofen were repeated once daily if any symptoms of pain were identified. Rats were allowed 3 days of recovery post-surgery. After surgery animals were housed individually to prevent injury or damage to the implants.

### Sound presentations

Pure tones and broadband noise (BBN) were generated digitally online. The word-like stimuli were loaded from pre-synthesized files. All sound generation was performed using Matlab (The Mathworks, Inc.). The digital signals were transduced to voltage signals by a sound card (M-16 AD, RME), attenuated (PA5, TDT), and played through a stereo power amplifier (SA1, TDT) and a free field speaker (MF1, TDT) that was placed above the experimental cage. For pure tones, 0 dB attenuation corresponded to a sound level of about 100 dB SPL throughout the frequency range of the word stimuli.

### Electrophysiological recordings

Recordings were performed using an AlphaLab SnR™ recording system (Alpha Omega Engineering, Nazareth, Israel) connected to a TBSI transmitter-receiver system for wireless recordings (Triangle BioSystems International, Durham, NC, USA). The 64-channel transmitter and the battery were mounted onto a custom-made interconnector with a battery holder (total weight of the interconnector with the transmitter and the battery was approximately 15 g). Before each recording session, the device was attached to the electrode array.

Each of the four recording sessions (two before and two after conditioning) started with a characterization of the response properties of the recording location. First, we recorded responses to broad-band noise (BBN) using a sequence of 280 BBN bursts with a duration of 200 ms, 10 ms linear onset and offset ramps, ISI (onset-to-onset) of 500 ms, and seven different attenuation levels (0-60 dB with 10 dB steps). Levels were presented pseudo-randomly so that each level was presented 40 times.

Responses to tones were collected using quasi-random frequency sequences of 370 pure tone bursts (50 ms, 5 ms rise/fall time; ISI of 500 ms) at 37 frequencies (1–64 kHz, 6 frequencies per octave). The sequences were presented at decreasing attenuation levels, starting at 10 dB attenuation with 10 dB steps until the threshold of the neural activity was reached (usually at 50-60 dB attenuation). On the first day of recording, these data were used to select the main frequencies and sound levels for all behavioral tests using tones. The best frequency (BF) was determined as the frequency that gave rise to the strongest responses in most electrodes. Two frequencies evoking large responses were selected on either side of the BF, symmetrically, for further study. The lower frequency was denoted f1, the higher was denoted f2, and they were selected such that f2/f1=1.44. We then recorded responses to oddball sequences consisting of the word stimuli and (separately) of pure tones of the two selected frequencies.

### Oddball sequences

Tone oddball sequences consisted of 30 ms (5 ms rise/fall time) pure tone bursts, presented with an interstimulus interval (ISI, onset to onset) of 300 ms. Each sequence contained 25 deviants and 475 standards in a pseudo-random order, so that the deviant frequency had a probability of 5%. These are the conditions used in most SSA studies coming from our lab (7,26). Two oddball sequences have been used, one with f1 standard and f2 deviant, and the other with the roles of the two frequencies reversed.

The word stimuli (‘danger’, phonetically ‘/deındƷər/’, and ‘safety’, phonetically /seıfti/, respectively) were computer generated by an open-source text to speech synthesizer (Festival, Linux, Fedora 14) and modified using the STRAIGHT vocoder (Kawahara et al. 2008) and Matlab routines. The frequency content of the two sounds was shifted above 1 kHz and the pitch contour was set to a constant 350 Hz in order to remove pitch cues for word identity. The total energy and power spectra of the two sounds were equalized in order to remove simple energy and spectral cues for word identity. These modifications resulted in sounds that had some features of speech, notably strong spectro-temporal modulations in the speech range (Supplementary Fig. 1). Oddball Sequences consisting of word stimuli were presented at a rate of 1 Hz. The deviant word (either “danger” or “safety”) had a probability of 5%, and the oddball sequences consisted of 500 stimuli (475 standards and 25 deviants). Two sequences were presented. In one sequence, the standard was “danger” and the deviant was “safety”. In opposite sequence “danger” was the deviant and “safety” the standard. presentations were counterbalanced.

### Fear conditioning

We used a discriminative fear conditioning protocol, loosely adapted from Letzkus et. al. (2011). The rats were exposed to 20 blocks of sounds, alternating between CS- and CS+, with silent intervals of 60 - 180 s (randomly selected) between the blocks (Fig. 1C). Each block consisted of a 30 s train of one of the stimuli at a sound pressure level of 70 dB. The CS+ was paired with a foot shock (2 s, 0.4 mA). The onset of the foot shock was 2 seconds before the end of the sequence.

Each word was used (in different groups of rats) as CS+ and as CS-. During the 30 s sequences, the stimuli were presented at 0.5 Hz (once every 2 s). A pseudo-conditioned group was subjected to the same procedure (using the word stimuli) but without applying foot shocks.

To condition with tones, the CS+ and CS- sequences consisted of 30 s sequences of pure tones of the two previously selected frequencies (30 ms tone pips, presented every 300 ms, 5 ms linear onset/offset ramps).

On the two days following conditioning, the rats were submitted to fear retrieval test in context B, during which they were exposed alternately to presentations of the CS– and of the CS+ sound sequences, for a total of 4 times each (Fig. 1D).

### Behavioral analysis

To determine the amount of freezing, we monitored rat behavior using a ceiling mounted CCD video camera (DFK 23G445, The imaging source, Taipei city, Taiwan). Video images (30 frames/s) were later analyzed and synchronized with behaviorally-relevant events (sound and shock presentations) using custom Matlab routines (supplementary Fig. 2). Each frame was smoothed with a Gaussian filter with a width of 10 pixels and transformed into grayscale. Each frame was subtracted from the previous one, the difference images were thresholded, and the number of non-zero pixels provided a measure of the amount of movement from one frame to the next. The amount of movement was smoothed over 2 s periods (boxcar smoothing, 59 points at 30 Hz), and freezing was detected when the smoothed trace decreased below a threshold. This procedure had two free parameters, the threshold for the detection of pixels that changed in the temporal difference images, and the threshold for detecting freezing. These were determined to fit best a set of test cases scored manually for the amount of freezing.

Mean freezing was calculated for 40 s following the beginning of each stimulus block (block duration + 10 s). Baseline freezing was calculated from the first two minutes of each session, before the presentation of the first stimulus block.

The amount of freezing in the different conditions was analyzed using a linear mixed effects model (Matlab, function fitlme). The fixed factors were the experimental group (conditioned to words, conditioned to tones, pseudo conditioned) and stimulus condition (Baseline, CS-, CS+), with rats within groups used as a random factor.

### Analysis of the electrophysiological data

The data were analyzed using Matlab. Local field potentials (LFPs) were extracted from the raw electrode signals by lowpass filtering (corner frequency: 200 Hz) and downsampling from 22 to 1 kHz.

For the tone responses, LFP responses were baseline corrected to the 50 ms before stimulus onset. The peak negative response was identified in the 40 ms time window starting at stimulus onset, and response strength was quantified by averaging the LFP over the 9 ms window centered on the peak.

For the word responses, LFP responses were baseline corrected to the 50 ms before onset of the first vowel (the justification for this procedure is described in the Results section). The peak negative response was found in the 40 ms time window starting at the onset of the first vowel, and response strength was quantified by averaging the LFP over the 9 ms window centered on the peak.

The responses to a given stimulus were included in the final dataset when there was a significant response in at least one of the conditions (standard, deviant, before conditioning, after conditioning). Significance test was performed by a paired t-test between the set of single-trial responses (same response window as above) and the corresponding pre-stimulus LFP (p<0.05).

In order to quantify the effect of probability on tone responses, the contrast between the responses to the same stimulus when it was standard and when it was deviant was used. This contrast is termed SSA index (SI, Ulanovsky et al. (1)):

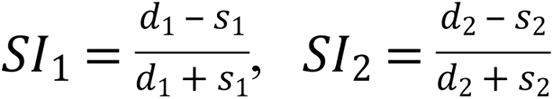

Where d_i_ and s_i_ represent the responses to the two different stimuli (i=1,2) when they were deviant and standard respectively.

The responses to the word stimuli were analyzed using a linear mixed effects model (Matlab routine fitlme). The fixed factors were the behavioral role of the sound (CS+/CS-, conditioned to tones, pseudo conditioned), sound probability (standard/deviant), and time (before/after conditioning). Stimulus type (’safety’ or ‘danger’), rat, recording session (1^st^ or 2^nd^), and electrode within rat were entered as random factors. The responses to the tones were analyzed using a similar model: behavioral role (CS+/CS-, conditioned to words, pseudo-conditioned), sound probability and time. Stimulus type (low or high frequency), rat, recording session (1^st^ or 2^nd^), and electrode within rat were entered as random factors. Table 1 reports the results for all fixed effects (Matlab routine anova). All main effects and almost all interactions were significant, often highly so. We therefore report later the results of post-hoc tests of individual contrasts between fixed effects (coefficient tests using the Matlab routine coefTest, performing an F test for the specific contrast against the null hypothesis that it is zero).

**Table 1.**

| **Responses to word-like stimuli** |  |  |
| --- | --- | --- |
| *Main effects* |  |  |
| Sound probability (standard/deviant) | F(1,3979)=5.3 | P=0.021 |
| Time (before/after conditioning) | F(1,3979)=6.2 | P=0.013 |
| Behavioral role (CS+,CS-,Pseudo-conditioning,conditioned to tones) | F(3,3979)=6.6 | P=2.0*10<sup>-4</sup> |
| *Interactions* |  |  |
| Probability x Time | F(1,3979)=16 | P=6.6*10<sup>-5</sup> |
| Probability x Behavioral role | F(3,3979)=5.9 | P=5.5*10<sup>-4</sup> |
| Time x Behavioral role | F(3,3979)=4.7 | P=2.7*10<sup>-3</sup> |
| Probability x Time x Behavioral role | F(3,3979)=11 | P=4.3*10<sup>-7</sup> |
| **Responses to Tones** |  |  |
| *Main effects* |  |  |
| Sound probability (standard/deviant) | F(1,3860)=75 | P=5.4*10<sup>-18</sup> |
| Time (before/after conditioning) | F(1,3860)=32 | P=2.0*10<sup>-8</sup> |
| Behavioral role (CS+,CS-,Pseudo-conditioning,conditioned to words) | F(3,3860)=3.0 | P=0.029 |
| *Interactions* |  |  |
| Probability x Time | F(1,3860)=0.92 | P=0.34 |
| Probability x Behavioral role | F(3,3860)=8.1 | P=2.1*10<sup>-5</sup> |
| Time x Behavioral role | F(3,3860)=7.5 | P=5.0*10<sup>-5</sup> |
| Probability x Time x Behavioral role | F(3,3860)=13 | P=3.7*10<sup>-8</sup> |

## Results

Twenty-one rats underwent the full experimental procedure (10 conditioned to words, of which 5 rats were conditioned to ‘safety’ and 5 rats to ‘danger’; 6 conditioned to tones; 5 pseudo-conditioned, of which only 4 have valid behavioral data). The behavioral results are summarized in Fig. 2.

**Fig 2.**
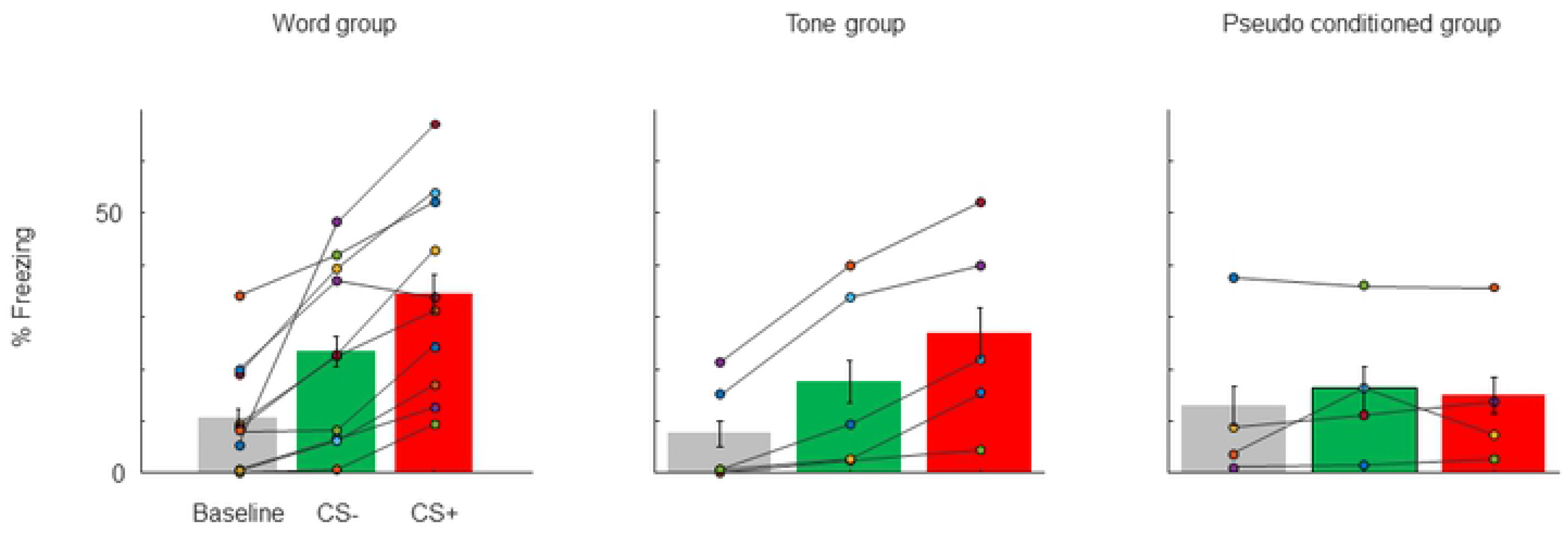
Behavioral results. Mean percentage of freezing for the conditioned animals at baseline, during CS+ presentations, and during CS- presentations. Left: Word group. Middle: Tone group. Right: Pseudo-conditioning group. Error bars are standard error of the mean amount of freezing within animal.

The significant main effect of stimulus condition (F(2,471)=8.4, P=2.5*10^-4^) confirmed that freezing differed for presentations of different stimuli (Baseline, CS- and CS+) with less freezing at Baseline than at both CS+ and CS-. The main effect of conditioning group was not significant (F(2,471)=0.19, P=0.83), but there was a significant interaction between stimuli and group (F(4,471)=3.2, P=0.013) demonstrating that following conditioning, the different groups (conditioned to tones, to words, and pseudo-conditioned) showed different patterns of freezing.

All rats in the word group froze more during CS+ than at baseline (Fig 2, left). There was also a generalization: during CS- presentations, all rats also showed elevated freezing relative to baseline. However, almost all rats (9/10) froze more when CS+ was presented than when CS- was presented. A post-hoc test showed a significant difference between freezing during CS+ and CS- presentations (F(1,471)=11, P=8.0*10^-4^).

The rats conditioned with tones (Fig. 2, middle) displayed a similar pattern: they froze during both CS+ and CS-, but more to CS+. In this group the difference between freezing for CS+ and CS- was not significant (F(1,471)=1.8, P=0.18). This could be due to the small number of animals used for this test, or to the small frequency interval between CS+ and CS- (half octave).

In the pseudo conditioned group (Fig 2, right; the behavioral data of one of the animals was not recorded) there was no significant increase in freezing for either stimulus relative to baseline (F(2,471)=0.33, P=0.72).

We recorded LFPs from 336 recording locations in 21 rats. Figure 3 shows the population averages of the responses to the two word stimuli. Since the word stimuli had a complex temporal structure, the responses included multiple temporal components. Figures 3a and 3b display, from top to bottom, the waveforms of the word stimuli, the average of all responses in all animals, and the responses in each individual animal averaged over all electrodes. For both words, the first response component was evoked by the onset of the initial consonants (/d/ and /s/), at 270 ms after trial onset for “danger” and 302 ms after trial onset for “safety”. The next response component, which was the largest one for both words, was evoked by the onset of the first vowels (/ei/), at 395 ms “danger” and 298 ms for “safety”. The third response component was evoked by the final consonant of the 1st syllable of each word (/n/ and /f/, 592 ms “danger”, 565 ms “safety”). The fourth component was evoked by the onset of the second vowel (/er/ and /i/, 690 ms “danger”, 630 ms “safety”).

**Fig 3.**
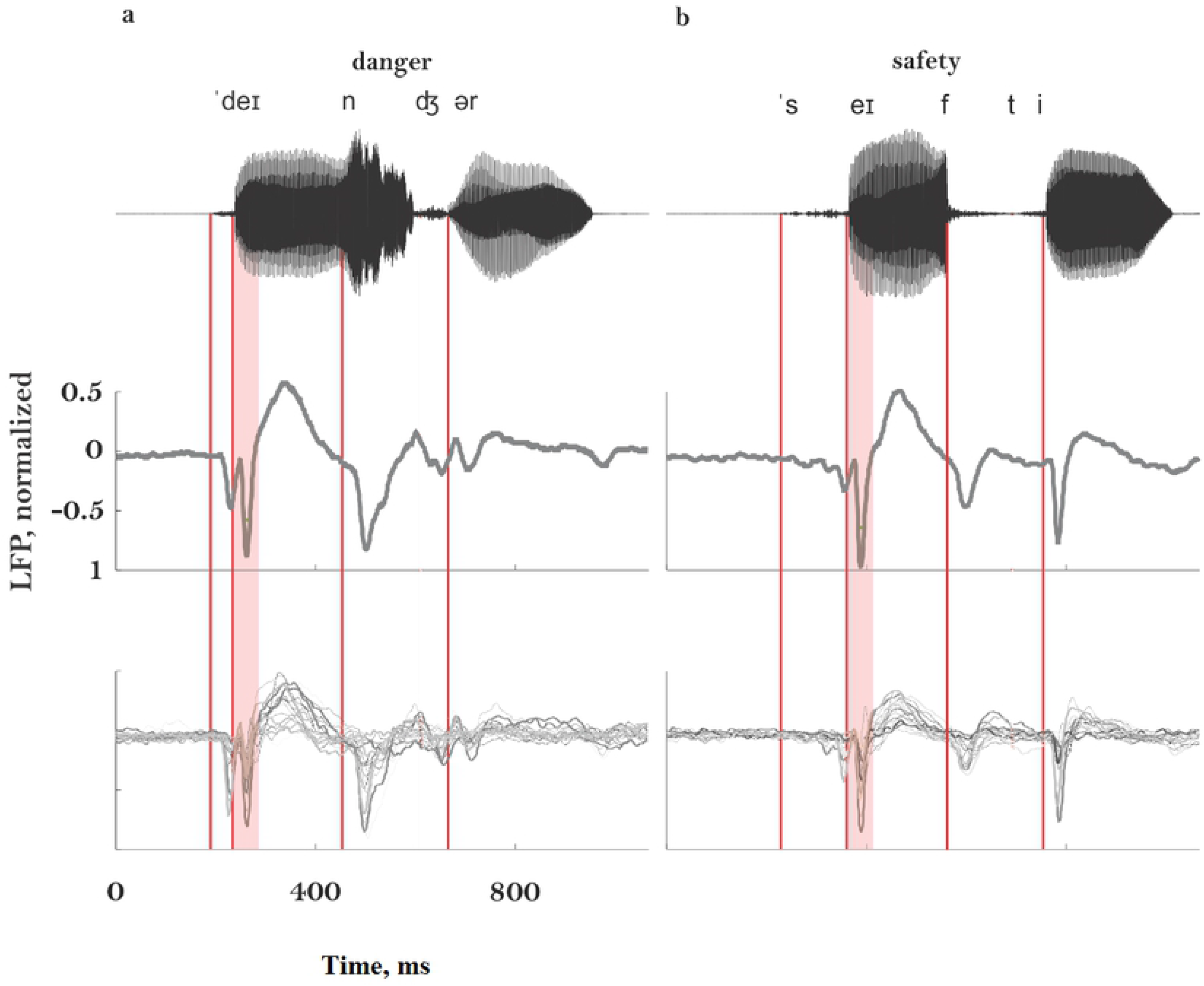
Responses to word stimuli. The waveform of the word stimuli (top), mean LFP responses over all electrodes and all animals (middle), and the average responses for all electrodes within animal, plotted for all animals (bottom). (a) For the word ‘danger’. (b) For the word ‘safety’.

As seen at the bottom of Fig. 3, response components showed high variability between animals. We therefore report all our results for the onset response to the first vowel of each word, a component that occurred in all animals. The responses at other time windows generally changed in parallel with these responses (27).

Responses to the word-like stimuli and to tones were collected in all three groups (word group, tone group, and the pseudo-conditioned group). All main effects and interactions were significant (Table 1), showing that SSA was present (main effect of probability both for words and for tones) and that conditioning indeed modified the responses in ways that depended on the probability as well as on the behavioral role of the stimulus. We therefore report below the results of post-hoc tests for the specific contrasts of interest.

We first discuss the effects of conditioning on SSA. Figure 4 summarizes these data. Each panel shows the distribution of the SSA indices computed before and after conditioning, for the word-like stimuli (top row) and the tone stimuli (bottom row). Control A consists of the SSA recorded in rats conditioned to the other stimulus (to tones for the word-like stimuli, to word-like stimuli for the tone stimuli). Control B consists of the recordings in the pseudo-conditioned rats.

**Figure 4.**
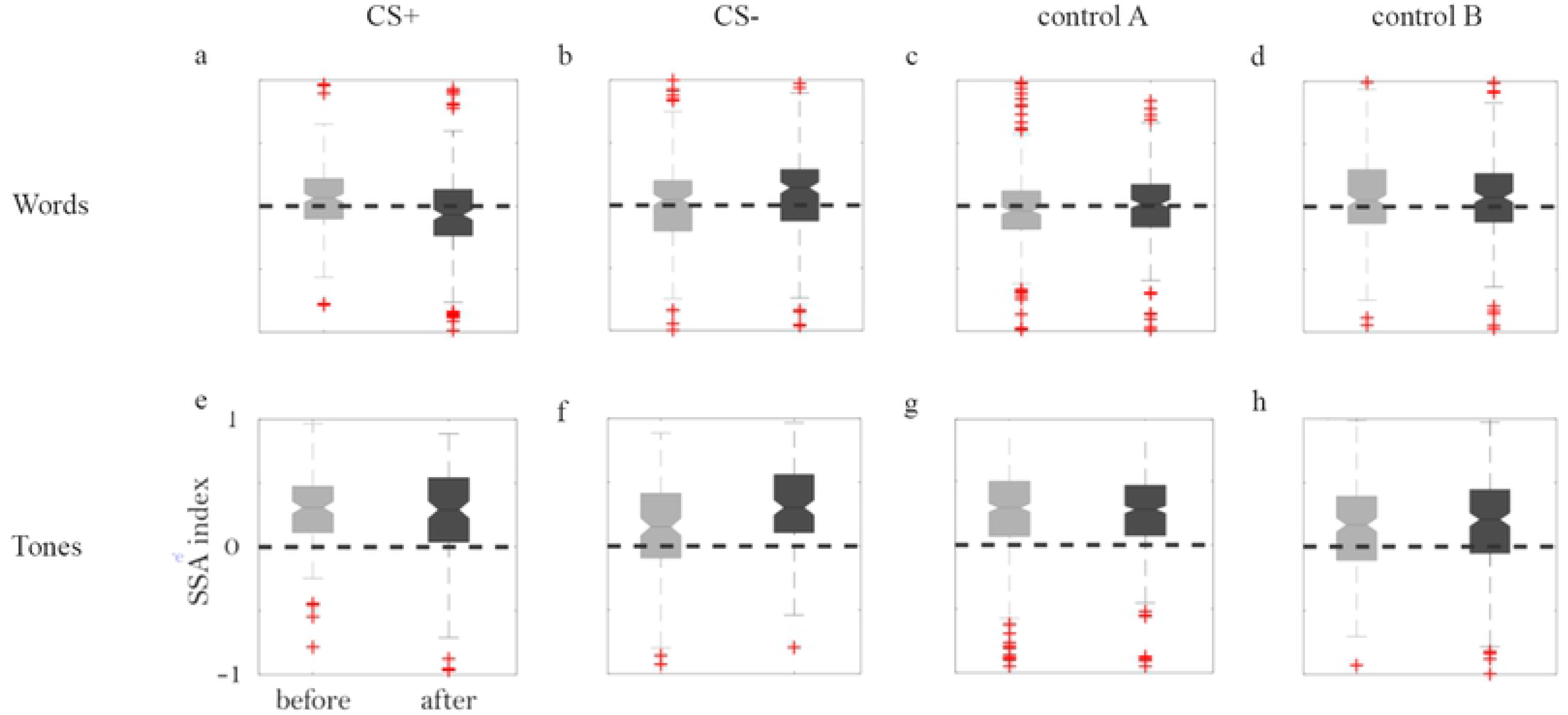
Changes in SSA following conditioning. (a) Box plots showing the distribution of SSA indices before (left, gray) and after (right, black) conditioning, for word stimuli used as CS+. The dashed line is at 0, corresponding to equal responses to standards and to deviants. The box is centered on the median of the distribution and its top and bottom edges indicate the 25^th^ and 75^th^ percentiles of the distribution. Outliers are marked by red plus signs, and the whiskers show the extent of all the data that is not considered as outliers. The notches represent 5% confidence intervals around the medians. (b) The same for word CS- stimuli. (c) The same for the responses to word stimuli recorded in animals conditioned with tones. (d) The same for responses to word stimuli recorded in pseudo-conditioned animals. (e)-(h) The same, for tone stimuli. In this case, control A consisted of recordings of tone responses in animals conditioned with word-like stimuli.

We hypothesized in the introduction that SSA would decrease for the CS+ stimuli and increase for the CS- stimuli. When the word-like stimuli served as CS+, there was a significant decrease of the SSA index following conditioning (Fig 4a, −13%, F(1,1808)=16, P=8.3*10^-5^). In contrast, when they served as CS-, there was a moderate but significant increase of the SSA index following conditioning (Fig 4b, 7%, F(1,1808)=5.1, P=0.025). In both control groups, the SSA index did not change significantly following conditioning (Fig 4c, control A, tone-conditioned animals: 12%, F(1,1808)=0.87, P=0.35; Fig. 4d, control B, pseudo-conditioned: 0.7%, F(1,1808)=0.054, P=0.82). Thus, the change in SSA was specific to the stimuli that gained behavioral meaning; the SSA decreased for the CS+ and increased for the CS- stimuli. These results are fully consistent with our hypothesis.

When tones served as CS+, the SSA index to remain largely the same after conditioning (Fig. 4e, −5%, F(1,1801)=1.5, P=0.2). For CS- tones, a highly significant increase in SSA occurred after conditioning (Fig 4f, 16%, F(1,1801)=18, P=2.3*10^-5^). In both control groups, there was virtually no change in SSA index after conditioning (Fig 4g, control A, word-conditioned group: −0.4%, F(1,1808)=0.023, P=0.88; Fig. 4h, control B, pseudo-conditioned group: 1%, F(1,1801) =0.14, P=0.7). This pattern was partially consistent with our working hypothesis. The change in SSA was specific to stimuli that gained behavioral meaning; the SSA to CS- stimuli increased, as hypothesized, while the SSA to the CS+ stimuli didn’t change significantly,

Next, we examined the patterns of changes in the responses to standards and deviants that underlay the changed SSA. SSA can change because the responses to standards changed and/or because the responses to deviants changed, and we wanted to find out which pattern actually occurred.

Figure 5a illustrates the most surprising finding. It displays the average responses to the word-like stimuli used as CS+ when they were tested as deviants, before (light red) and after (dark red) conditioning. The responses showed a substantial and highly significant decrease, rather than the expected increase (Fig 5c, deviants: −43%, F(1,3979)=66, P=5.6*10^-16^). The responses to standards also decreased significantly, although to a lesser degree, following conditioning (Fig 5b and 5c, standards: −15%, F(1,3979)=6.2, P=0.013). The significant decrease in SSA shown by the word-like stimuli used as CS+ (Fig. 4a) can be traced therefore to the fact that following conditioning, the responses to the CS+ word-like stimuli, when used as deviants, decreased more than the responses to the same stimuli when used as standards.

**Figure 5.**
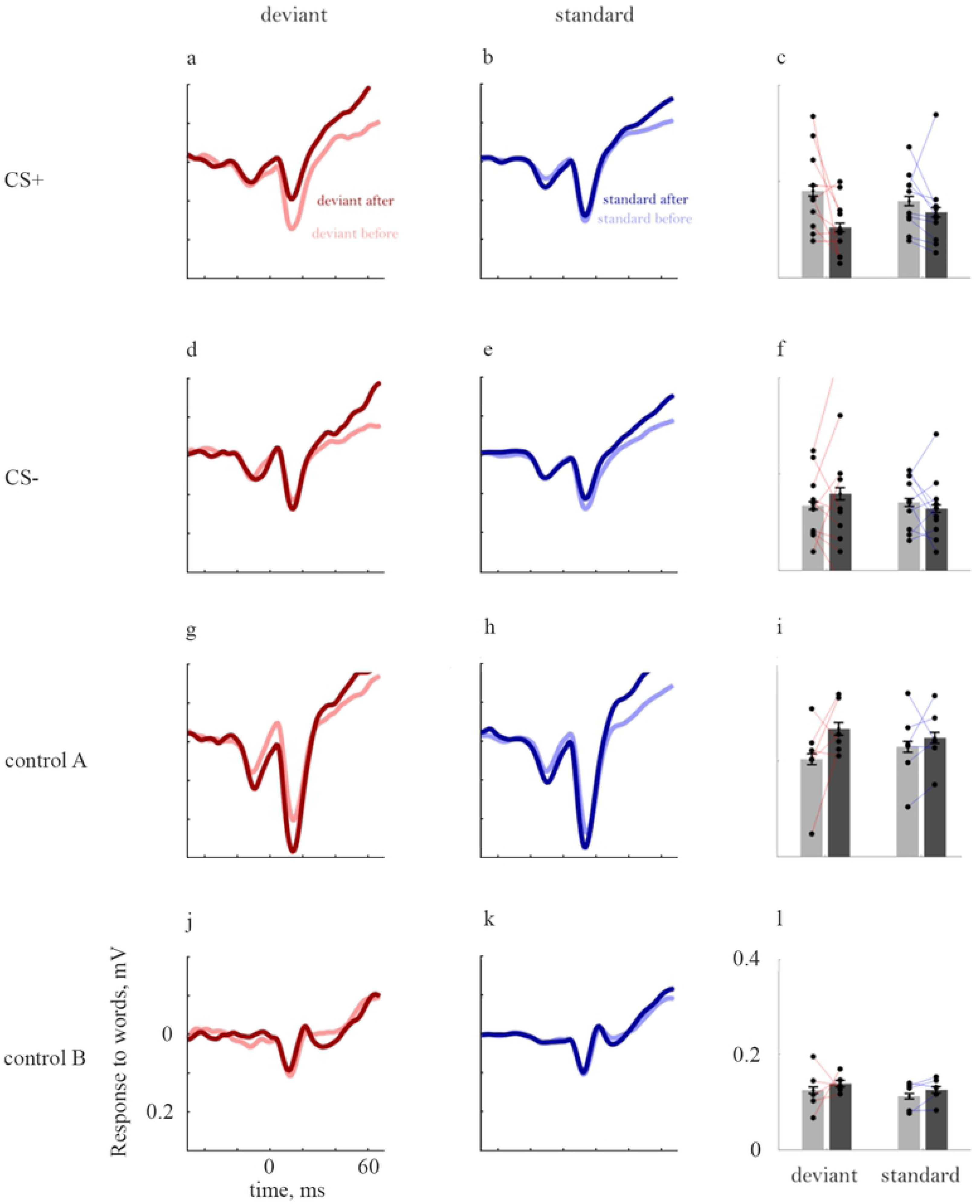
Changes in the responses to word-like stimuli following conditioning. (a) Responses to CS+ word-like stimuli when deviants. The light red shows the responses when these stimuli were tested as deviants before conditioning. During conditioning, the same word-like stimuli were used as CS+. The dark red line shows the responses to the same stimuli when tested again as deviants after conditioning. The time window shown starts at stimulus onset and ends 40 ms after the onset of the first vowel. (b) The same, for the responses to the CS+ word stimuli when tested as standards. (c) Average peak responses to the same stimuli. The bars represent the average peak response before (gray) and after (black) conditioning. The dots represent the average peak response across all electrodes and sessions in each animal. The peak responses before and after conditioning in each individual animal are connected with a line. (d-f) The same, for word-like CS- stimuli. (g-i) The same, for the responses to word-like stimuli tested in rats conditioned with pure tones. (j-l) The same, for the responses to word-like stimuli tested in pseudo-conditioned rats. Note that in this case, while the average response (panel j) was slightly smaller after than before conditioning, the average peak response (panel l) was slightly larger. The reason for such discrepancies here and elsewhere is the fact that peak responses were determined in each electrode and animal individually, and therefore could occur at time points that are different than the time point of the peak response following averaging.

This decrease in the responses following conditioning was restricted to the word-like stimuli when used as CS+ during conditioning. Indeed, when used as CS-, following conditioning the responses to word-like stimuli increased when deviant (Fig 5d and 5f, deviants: 19%, F(1,3979)=7.2, P=0.0074) and did not change significantly when used as standards (Fig 7e and 7f, standards:, −6%, F(1,3979)=0.77, P=0.38). When tested in animals that have been conditioned to tones (control A, Figs. 5g-i), the responses to the word-like stimuli increased (deviants: 29%, F(1,3979)=49, P=3.0*10^-12^, standards: 8.2 %, coefficient test: F(1,3979)=5.1, P=0.024; Fig 6i-k). The responses to the word-like stimuli did not change significantly in the pseudo-conditioned animals (deviants: 12%, coefficient test: F(1,3979)=2.4, P=0.12; standards: 11%, coefficient test: F(1,3979)=2.3, P=0.13; Fig 5j-l).

**Figure 6.**
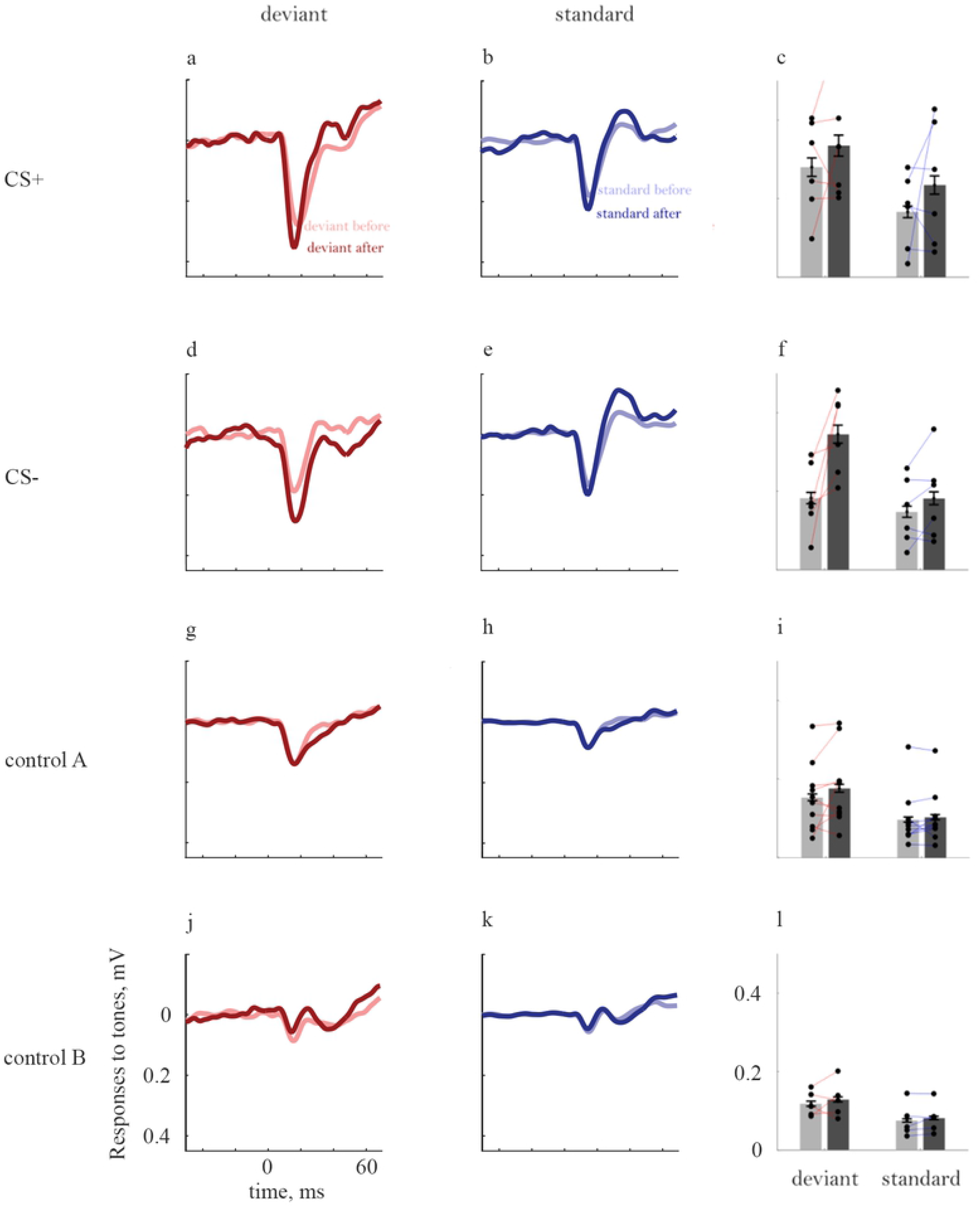
Changes in the responses to tones following conditioning. Same conventions and structure as Fig. 5. Control A (g-i) in this case consists of the responses to tone stimuli in animals that have been conditioned to word-like stimuli.

Responses to tones (Figure 6) showed, if anything, only increases following conditioning, as expected (28). Responses to CS+ tones increased following conditioning both when deviant (Fig 6a and 6c, deviants: 20%, F(1,3860)=18, P=2.1*10^-5^) and when standards (Fig 6b and 6c, standards: 45%, F(1,3860)=32, P=2.0*10^-8^). The Responses to CS- tones also increased, both when deviants (Fig 6d and 6f, deviants: 94%, coefficient test: F(1,3860)=149, P=1.0*10^-33^) and when standards (Figs 6e and 6f, standards: 25%, coefficient test: F(1,3860)=6.7, P=9.8*10^-3^).

The responses to deviant tones presented to rats conditioned to word-like stimuli increased significantly after conditioning (Fig 6g and 6i, deviants: 16%, F(1,3860)=9.9, P=1.7*10^-3^), while the responses to the same stimuli when standards in animals conditioned to the word-like stimuli did not change significantly (Fig 6h and 6i, standards: 7.9%, F(1,3860)=0.87, P=0.35). Thus, the decrease in the responses to word-like stimuli used as CS+ did not generalize to other stimuli in these rats. Responses to tones didn’t change significantly in pseudo-conditioned animals (deviants: 7.5%, coefficient test: F(1,3860)=0.90, P=0.34; standards: 8.8%, coefficient test: F(1,3979)=0.27, P=0.60; Figs. 6j-l).

## Discussion

We used fear conditioning to assign a behavioral meaning to complex sounds and to pure tones and then measured SSA elicited by these sounds before and after conditioning.

### Methodological issues

The current study was designed with the goal of recording neural signals in the same awake animals before and after conditioning, in order to allow within-animal comparison of the resulting electrophysiological changes. This experimental design made the study statistically powerful, but resulted in a long-duration protocol that made it difficult to collect stable spiking activity. Thus, the paper is based on recordings of LFPs.

LFPs are useful indices of neuronal activity, but need to be interpreted carefully. LFP measures the total synaptic input (rather than spiking output) near the electrode tip (29). LFPs integrate currents over relatively long distances – at least 1 mm (30) - and are therefore less local than recordings of spiking activity (31). Nevertheless, LFPs are often interpreted as an index of spiking activity. Indeed, there are many experimental observations showing correlated changes in the two signals (32–34), including in auditory cortex. These correlations presumably have to do with the fact that most of the input currents in cortex are produced by local sources and therefore correlate with the overall spiking activity. Given the many demonstrations of such a correlation in auditory cortex, we accept it for the rest of the discussion.

### Conditioning differentially affected SSA to behaviorally meaningful sounds

Here we used the powerful classical fear conditioning paradigm in order to assign two possible meanings to sounds: a sound could either predict an aversive consequence (CS+) or predict the lack of an aversive consequence (CS-). CS- sounds are behaviorally meaningful – they occurred in 50% of the trials, and informed the rat that a shock was not imminent. Thus, we expected changes in SSA to occur for both types of sounds. In addition, we tested SSA using sounds that have not been used in the conditioning session (tones for the rats conditioned with words, and words for the rats conditioned with tones).

Our working hypothesis suggested that SSA to CS+ sounds should decrease and SSA to CS- sounds should increase, while SSA to sounds that have not been used during the conditioning session should be mostly unaffected. Our results are largely consistent with this hypothesis: at least at the population level, SSA was affected by conditioning as expected from functional considerations – following conditioning, responses to CS+ stimuli adapted to a similar degree or less, while responses to CS- stimuli adapted more than before conditioning.

### Conditioning differentially affected responses to tones and to complex sounds

Although the changes in SSA roughly followed our working hypothesis for both tones and word-like stimuli, the changes in response strengths that underlay the changes in SSA showed an unexpected pattern. While response strength generally increased when the conditioned stimuli were tones, response strength to word-like CS+ stimuli decreased substantially and consistently following conditioning.

Fear conditioning has been almost invariably associated with increased responses to the CS+ stimulus in auditory cortex (24,23). In the experiments described here, the ubiquitous findings of increased responses to CS+ stimuli were reproduced for the tone stimuli. In fact, in animals conditioned to tones, responses to both CS+ and CS- tones, as well as to the word-like stimuli, all increased following conditioning.

For the word-like stimuli, on the other hand, conditioning affected differentially the size of the responses to CS+ and CS- stimuli. Responses to word-like stimuli when CS+ showed an unexpected decrease. This decrease was specific to the behavioral role of the stimulus: responses to the CS- word tended to increase when deviant and showed a non-significant decrease when standard. The decrease was also specific to the acoustic nature of the stimuli: in the same animals, responses to tones increased moderately following conditioning.

The specific decrease in the responses to word-like CS+ stimuli is one of the largest effects in this study. It occurred when the CS+ word was tested as deviant as well as when standard. Since deviant responses decreased substantially more than standard responses, SSA decreased significantly following conditioning. In fact, the SSA index became negative on average: responses to repeated CS+ stimuli were on average somewhat larger than to rare ones.

To the best of our knowledge, previous research has shown two exceptions to the ubiquitous increase in the responses to the CS+ stimuli. The first is plasticity in the highly specialized auditory system of the Jamaican mustached bat, *pteronotus parnellii*, evoked by microstimulation of auditory cortex. Following this manipulation, the neurons in the stimulated region showed shifts of their frequency tuning away from the characteristic parameters of the stimulated point (23). Such shifts have been observed throughout the auditory system (in cortex, auditory thalamus and inferior colliculus) when microstimulation was performed in auditory cortex areas that were specialized for the processing of the echolocation calls (the DSCF area, the highly expanded area representing the 60 kHz component of the echolocation call, and the FM-FM area). Similar microstimulation experiments in non-specialized parts of auditory cortex gave rise to the expected tuning shifts towards the characteristic parameters of the stimulated area (35). Suga and his colleagues concluded the shifts of sensitivity away from those of the stimulated area is a property of the specialized processing areas in the bat auditory cortex (23).

The current results with the word-like stimuli are reminiscent of this thread of results. Instead of shifting the responses towards the CS+, there is a shift of the responses away – reduction of the CS+ responses together with a potentially moderate increase in the responses to CS- stimuli as well as to tones. In contrast with the results of Suga and colleagues, we observed these shifts in an animal that is not an auditory specialist. Nevertheless, there is an interesting analogy – the ‘centrifugal’ (36) shifts in our experiments were observed only for complex stimuli that presumably engaged large territories of auditory cortex. We therefore suggest a possible reinterpretation of the observations of Suga and his colleagues – it is not the difference between specialized and non-specialized processing, but rather the difference between the extent of cortex that is activated by the conditioned stimuli, that is responsible for the different patterns of results.

The second report of decreased responses to CS+ stimuli concerns operant conditioning experiments in ferrets (37). In animals trained to stop licking at target presentation, the responses to the target increased during task performance. In contrast, in animals trained to lick during target presentation, the responses to the target decreased during task performance. David and Colleagues (37) interpreted these results in terms of increased contrast between the target and non-target stimuli, in either case the larger responses being elicited by the stimuli that were associated with the aversive outcomes.

In the results reported here, increased and decreased responses to CS+ stimuli could be elicited independent of the behavioral paradigm, which was identical for all animals. Thus, both increased and decreased responses were associated with an aversive target (the CS+ stimulus), depending on whether it was narrowband (a pure tone) or wideband (a word-like stimulus). While there are substantial differences between our experiments and those of David et al. (37), at the least our results disprove a simple association of the polarity of response change with reward and punishment.

### Potential mechanisms

We interpret the changes in LFP as reflecting a corresponding change in the size of the spiking responses of the neuronal population around the recording electrodes. Given this assumption, our results provide two major constraints on mechanisms underlying these changes.

First, the changes documented here were a consequence of the conditioning procedure. This follows from the finding that neither SSA nor response strength changed significantly in the pseudo-conditioned rats. Thus, the plastic changes were initiated by the conjunction of cues that occur during the conditioning session, including the sounds and the aversive foot shocks. However, changes occurred also to the SSA evoked by CS- sounds, and in opposite direction to that evoked by CS+ sounds. Thus, plasticity occurred also in responses to sounds that were not directly associated with the aversive event, and even to sounds that were not presented at all during the conditioning sessions (tones in rats conditioned to word-like stimuli and word-like stimuli in rats conditioned to tones).

Second, the direction of the changes in response strength varied between tones and word-like stimuli. Responses to sounds in rats conditioned to tones increased to all stimuli (tones used as CS+, tones used as CS-, and word-like stimuli that were not used during conditioning). In contrast, the responses in rats conditioned to words specifically decreased to words used as CS+, while increasing somewhat to words used as CS- as well as to tones.

One mechanism that has been suggested to increase the responses to important sounds is increase in the release probability of glutamate, either at the thalamo-cortical or at the cortico-cortical synapses. In this case, deviant responses are expected to increase, but the increased synaptic depression consequent on the increased transmitter release is expected to decrease standard responses, leading to larger SSA. Such an effect has been demonstrated in consequence to environmental enrichment (38) – responses to sounds increased, but so did paired-pulse depression. The effects of conditioning on the responses to CS- stimuli were consistent with this mechanism. In rats conditioned to tones, the responses to both deviants and standards CS- tones increased, with larger increases of the deviant responses. In rats conditioned to word-like stimuli, the responses to deviant CS- stimuli increased while the responses to the same stimuli when standards did not change significantly.

On the other hand, responses to CS+ tones increased, but the SSA index did not change; and responses to word-like CS+ stimuli decreased in size and showed smaller (actually negative) SSA. All of these observations are inconsistent with simple increase in transmitter release probability.

The unexpected reduction of responses to word-like CS+ stimuli following conditioning could result from decreased excitation or from increased inhibition (or both). It is unlikely that excitation was greatly reduced, since responses to other stimuli (word-like CS- when deviant as well as to tones) were actually enhanced (admittedly, not by much). Thus, the main cause of the reduction in responses is most probably an increased inhibition evoked by the CS+ word-like stimuli.

Inhibitory effects may increase when excitatory-to-inhibitory synapses are potentiated, or when the inhibitory synapses themselves become more potent. Increased inhibition may then reduce the sensory responses to the CS+ stimuli. The reason inhibition would be potentiated more than excitation with word-like CS+ is unclear, but could be related to the large range of frequencies that were presumably affected during conditioning. For example, PV+ interneurons have wider tuning curves than nearby excitatory neurons (40). The use of broadband CS+ stimuli could potentiate more of the excitatory inputs to PV+ interneurons than the use of a pure tone CS+, leading to an overall greater inhibition (as in (36)).

### Conclusions

The results shown here demonstrate that SSA is shaped by experience. Whether a sound was used as CS+ or a CS- affected the subsequent degree of SSA it evokes. This finding resolves the main question that led to this study. At the same time, the use of complex sounds (word-like stimuli) led to the unexpected observation that responses to CS+ stimuli may actually decrease – even in the auditory cortex of a non-specialized mammal such as the rat, and even in the context of aversive conditioning. This finding suggests that the current understanding of plastic changes induced by a behavioral manipulation as simple as classical fear conditioning is still incomplete.

## Acknowledgements

This research was supported by an advanced ERC grant (project RATLAND, no. 340063), by F.I.R.S.T grant no. 1075/2013, and by ISF personal grant no. 390/2013 to IN, as well as by a grant from the Gatsby Charitable Foundation.

---

**Supplementary Figure 1.**
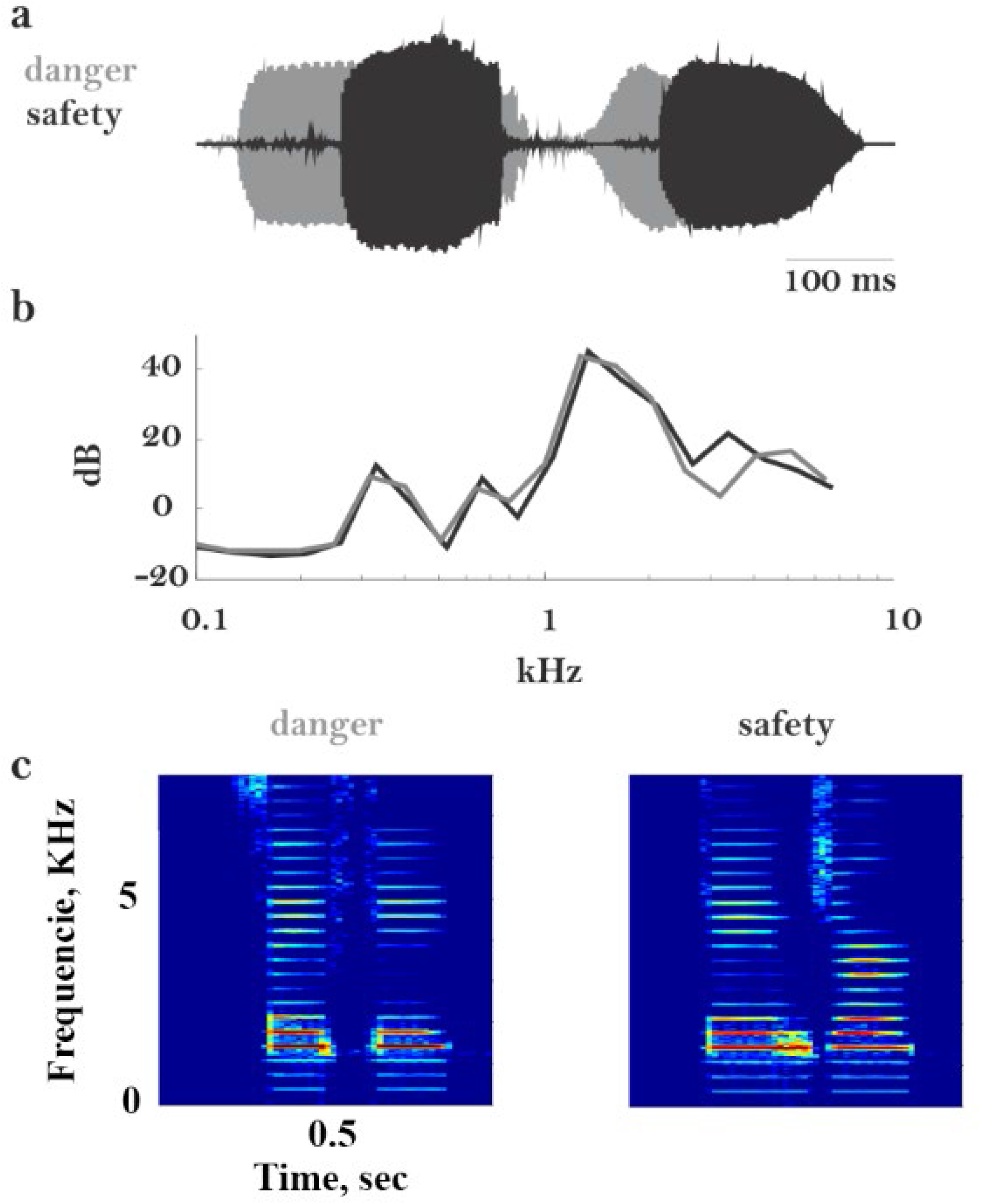
The word-like stimuli. Three different characterizations are displayed. a. Oscillograms of the two words. The two high-amplitude vowels in each word are clearly visible. b. Average power spectrum in 1/3 octave bands. The average power spectrum has been carefully equalized between the two stimuli. c. Spectrograms of the two stimuli. The ladder-like structures are the harmonics of the pitch of the two vowels, set to 300 Hz.

**Supplementary Figure 2.**
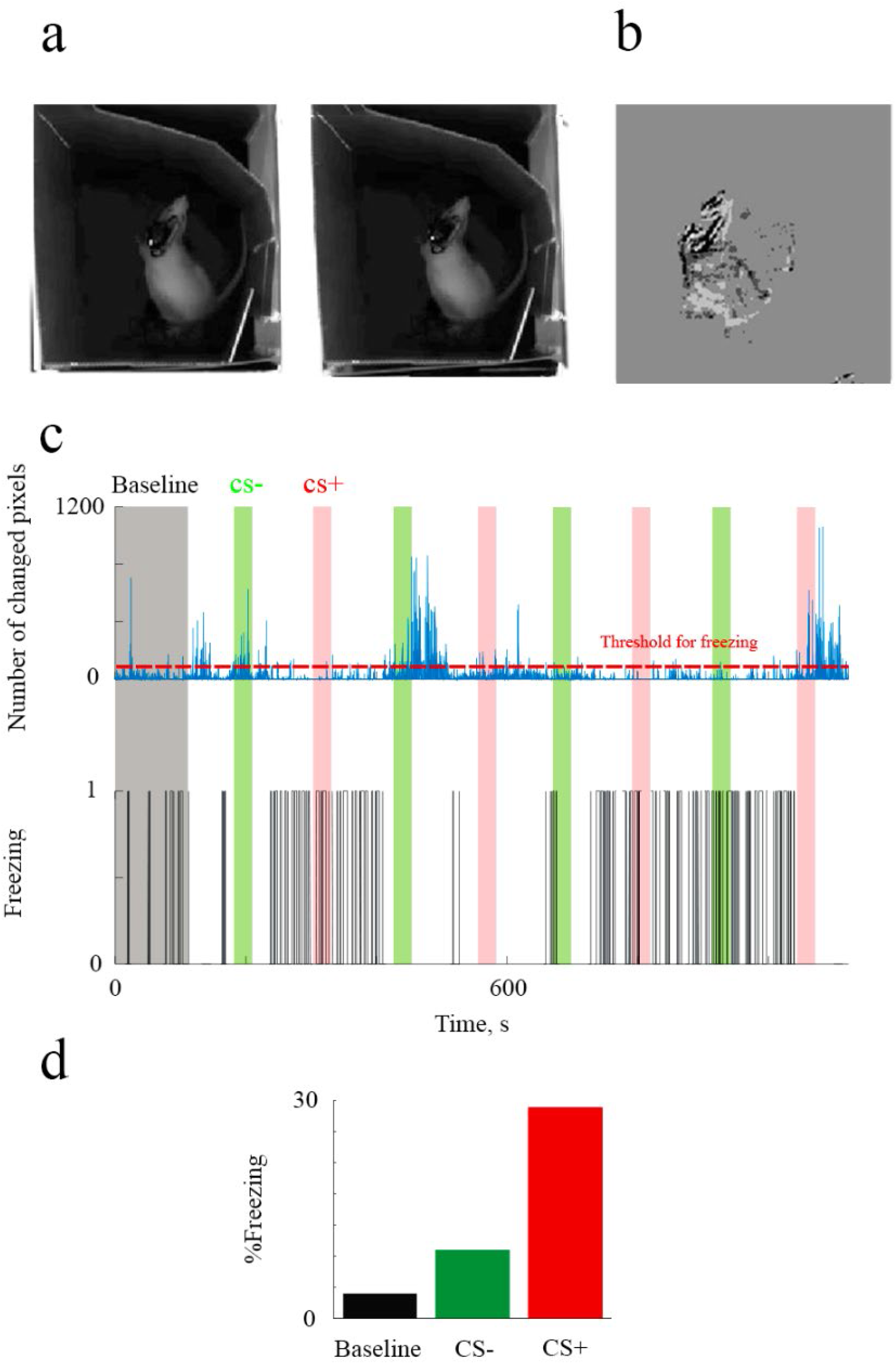
Detection of freezing. The algorithm used the video movie of the test episode. Starting from the individual frames (a), pairwise differences between successive frames were computed (b). The number of non-zero pixels was counted and smoothed. Panel c (blue) shows an example of such a smoothed trace. Freezing periods were determined by thresholding this trace (red line). Panel c (black) shows the resulting decisions. The fraction of time that freezing episodes occupied was determined separately for a baseline period (gray rectangle), for presentations of the CS+ (red) and for presentations of the CS- (green), as shown in panel d.

## Notes

#### Summary of Updates

Reordered figures; added supplemental figures to the file

